# An overview of data-driven HADDOCK strategies in CAPRI rounds 38-45

**DOI:** 10.1101/718122

**Authors:** P.I. Koukos, J. Roel-Touris, F. Ambrosetti, C. Geng, J. Schaarschmidt, M.E. Trellet, A.S.J. Melquiond, L.C. Xue, R.V. Honorato, I. Moreira, Z. Kurkcuoglu, A. Vangone, A.M.J.J. Bonvin

**Affiliations:** Computational Structural Biology Group, Department of Chemistry, Faculty of Science, Utrecht University, 3584CH, Utrecht, The Netherlands; Multiscale Materials Modelling and Virtual Design, Institute of Nanotechnology, Karlsruhe Institute of Technology (KIT), 76 021, Karlsruhe, Germany; CNC - Center for Neuroscience and Cell Biology; Rua Larga, FMUC, Polo I, 1°andar, Universidade de Coimbra, 3004-517; Coimbra, Portugal; PharmaResearch and Early Development, Large Molecule Research, Roche Innovation Center Munich, No nnenwald 2. 82377 Penzberg, Germany; Department of Physics, Sapienza University, Piazzale Aldo Moro 5, 00184, Rome, Italy

**Keywords:** Biomolecular interactions, complexes, integrative modelling, prediction, scoring

## Abstract

Our information-driven docking approach HADDOCK has demonstrated a sustained performance since the start of its participation to CAPRI. This is due, in part, to its ability to integrate data into the modelling process, and to the robustness of its scoring function. We participated in CAPRI both as server and as manual predictors.

In CAPRI rounds 38-45, we have used various strategies depending on the information at hand. These ranged from imposing restraints to a few residues identified from literature as being important for the interaction, to binding pockets identified from homologous complexes or template-based refinement / CA-CA restraint-guided docking from identified templates. When relevant, symmetry restraints were used to limit the conformational sampling. We also tested for a large decamer target a new implementation of the MARTINI coarse-grained force field in HADDOCK. Overall in the current rounds, we obtained acceptable or better predictions for 13 and 11 server and manual submissions, respectively, out of the 22 interfaces. Our server performance (acceptable models) was better (59%) than the manual (50%) one, in which we typically experiment with various combinations of protocols and data sources. Again, our simple scoring function based on a linear combination of intermolecular van der Waals and electrostatic energies and an empirical desolvation term demonstrated a good performance in the scoring experiment with a 63% success rate across all 22 interfaces.

An analysis of model quality indicates that, while we are consistently performing well in generating acceptable models, there is room for improvement for generating/identifying higher quality models.

## Introduction

In recent years molecular simulation techniques have been gaining traction as alternative methods for the elucidation of structural details underpinning cellular mechanisms^1^. Methods like molecular docking can complement experimental techniques such as X-RAY crystallography, NMR and cryo-electron microscopy and allow us to gain insight into the structural machinery of the cell. Understanding the structural elements at the core of interacting biomolecules in atomic detail is the first step toward understanding the nature of those interactions and being able to mechanistically explain what goes awry when they are disturbed (for example in disease phenotypes)^2–4^.

Initiatives like those of the D3R consortium^5,6^ test the ability of various modelling software to predict 3D structure and relative binding affinities of pharmaceutically interesting protein receptors bound to drug-like molecules. For protein-protein complexes, and to a lesser extent protein-peptide ones, the performance of various docking codes has been continuously evaluated over a period spanning almost 20 years in the worldwide CAPRI experiment^7–13^. We participated in all three experiments (server, manual and scoring) for all targets of rounds 38-45 with HADDOCK – our data-driven integrative modelling platform^14,15^. HADDOCK (High Ambiguity Driven DOCKing) makes use of biochemical/biophysical experimental information which is translated into distance restraints that drive the docking toward conformations that satisfy the experimentally available data. This cuts down on the need to exhaustively sample the conformational space and instead allows focusing on flexibly refining a subset of the models generated. Data-driven approaches have, of course, downsides as well, most prominently the fact that if the information that is provided to HADDOCK is incorrect, it will likely not sample the region of the conformational landscape close to the native state.

Our strategies for rounds 38-45 of CAPRI can be grouped in four categories depending on the nature of the target and the availability of information we could use to drive the docking. The first category concerns targets 121, 134 and 135 which featured protein-peptide complexes. Our approach for these targets boils down to variations of our protein-peptide protocol^16^ or threading on available templates and refining. The second category concerns targets 122 and 136 for which we could identify good template structures in the PDB^17^. Our main strategy here was to either superimpose the models we generated on the template structures and refine them or extract Cα-Cα interface distance restraints^18^ from the available templates, map them to the target sequence numbering and use those to drive the docking. The third – and most populated category – concerns targets for which no good templates were available but some experimental information was, mainly in the form of evolutionarily conserved residues or mutagenesis data. Targets 123-125 and 131-133 were modelled in this way. The last category concerns the protein-saccharide complexes (targets 126-130). For these targets we identified structures of the receptors of interest bound to oligosaccharide molecules of varying lengths and extracted binding site information in the form of the hydrogen bonds between the sugars and the residues of the pocket. We then used those restraints to dock conformers of the sugars we had generated from their SMILES strings.

In the following we describe the various strategies in more details, and present and discuss our results in light of the official CAPRI evaluation results, since not all reference structures of the complexes are yet available.

## Methods

Here we summarise the approaches we followed for the various targets of CAPRI rounds 38-45. Due to the diverse nature of the targets it is impossible to describe details of our protocols that hold true for all of them. Instead, we will first describe our high-level strategy and then list the details on a per-target basis. Additional details about the nature of the targets and the modelling process for all can be found in the SI.

Owing to the data-driven nature of HADDOCK, our first step after receiving the target sequences was to search structural databases like the PDB as well as make use of tools like HHPRED^19^ to identify close and remote structural homologues, respectively. Whenever high-quality templates that necessitated no mutations relative to our target sequences were available in the PDB we used those for the docking. Often though, differences in the sequences between the identified template structures and our targets meant that we needed to model the target sequence onto the template structure and to that end we used MODELLER^20^ and ROSETTA^21^. If we could not identify any usable templates then we turned to protein structure prediction servers such as RAPTOR-X^22,23^ or iTasser^24–26^.

After identifying or generating input models for the docking we searched the literature for putative interface information that might be available from (among others) bioinformatics predictions, mutagenesis or NMR titration experiments or even homologous complexes. For the targets where we could not identify any experimental information to assist in the modelling process, we made use of *ab initio* docking mode of HADDOCK, which defines restraints between the centres of mass of the various molecules and identified from an analysis of all generated models the most contacted residues.

After assembling all input structures and identifying all relevant interface information, we docked the individual molecules using HADDOCK. A typical HADDOCK run consists of the following three stages:

1. Random orientation of all molecules followed by energy minimisation-driven rigid body docking – it0.
2. Semi-flexible refinement in torsion angle space during which flexibility is introduced to the system starting with the interface (defined at a 5Å cut-off) sidechains before expanding to the interface backbone atoms as well – it1.
3. Short flexible refinement in explicit solvent – itw.

After every stage, the generated models are scored with the simple yet robust HADDOCK scoring function (HS) ^27,28^ which is a linear weighted sum of energetic and structural terms:

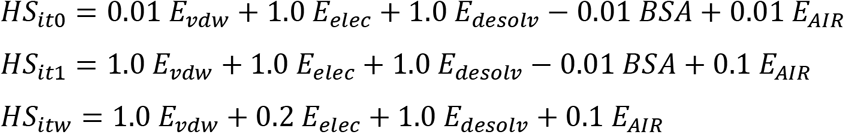

where *E_vdw_*, *E_elec_*, *E_desolv_* and *E_AIR_* stand for van der Waals, Coulomb electrostatics, desolvation and restraint energies respectively, and BSA for Buried Surface Area upon complex formation (in Å^2^). The non-bonded components of the score (*E_vdw_*, *E_elec_*) were calculated with the OPLS forcefield^29^, the desolvation energy is a solvent accessible surface area-dependent empirical term^30^ which estimates the energetic gain or penalty of burying specific sidechains upon complex formation.

For our participation in the server category, all models came from a single run that was submitted to the HADDOCK2.2 webserver, whereas for the manual category our submission usually consisted of a combined analysis of the various strategies that we tried out for each individual target. That combined analysis consists of clustering of all generated models using the Fraction of Common Contacts (FCC) metric with a cut-off of 0.75, meaning all models for which 75% of their interface residue contacts are shared would end up in the same cluster. The models are then scored according to the *HS_itw_* function shown above (minus the restraint term). The top clusters and models are then visually inspected, and the top 10 models that make up the submission are selected, typically selecting more models from the top ranked cluster and then spreading the remaining models over other good ranking clusters. Figure 1 provides a visual summary of the strategies we followed for some of our successful predictions.

**Figure 1:**
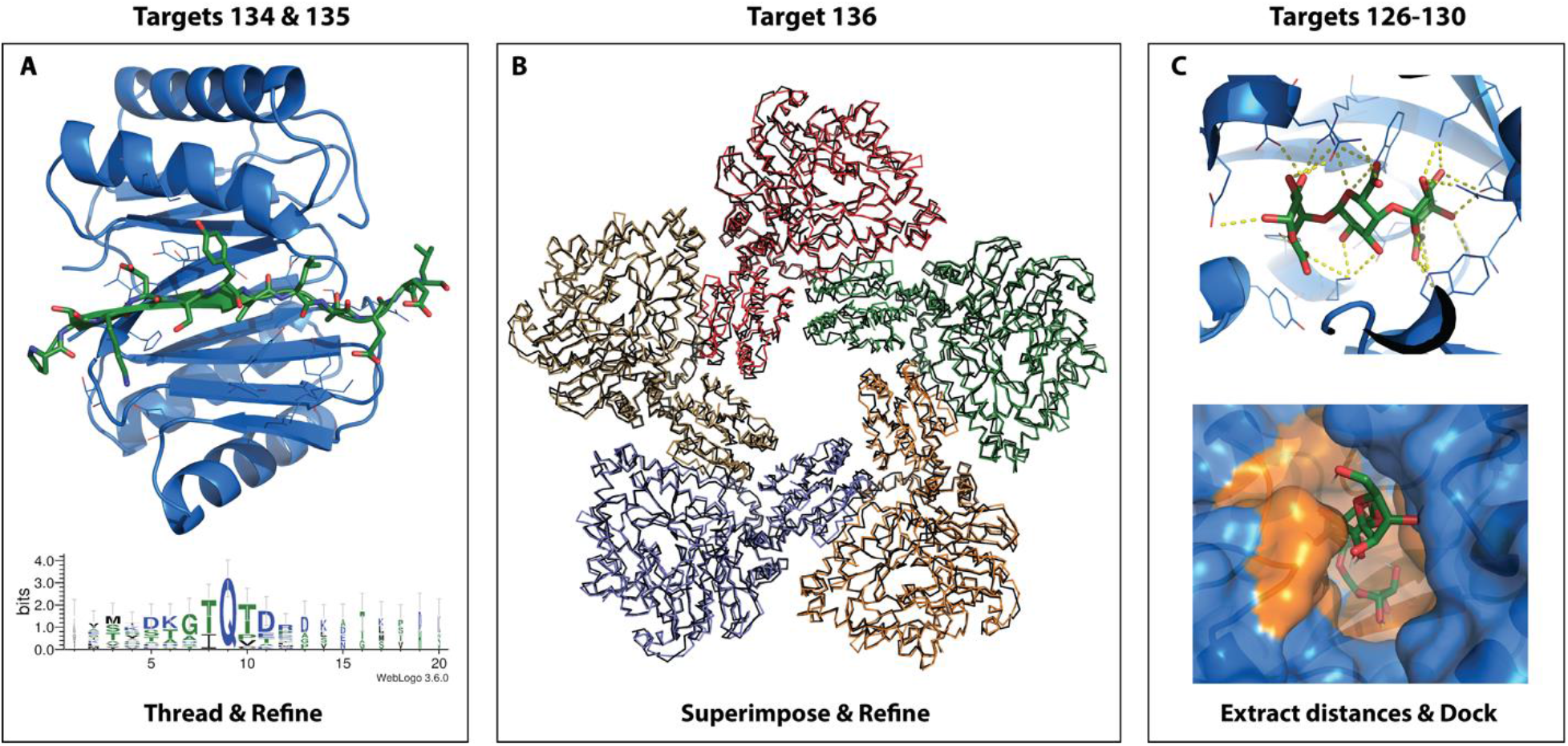
Visual summary for some of our successful predictions. Panel A concerns the strategy for the peptide targets, and specifically targets 134 and 135, which centred on threading and refinement. Our top–ranking model for the manual submission (medium quality) is shown. The receptor is depicted as cartoon and coloured blue with the binding site residues shown as lines, whereas the peptide is depicted as cartoon and sticks and coloured dark green. The bottom figure shows a sequence logo of all the peptides we identified in the literature which highlights a conserved TQT motif. Panel B concerns target 136 for which we generated models based on available templates, superimposed the models on respective templates and refined them. It depicts the dimer models we generated based on template 2vyc (only the top half of the decameric ring is shown to simplify the figure), superimposed on the template structure chains and refined in HADDOCK. The template structure is shown as black ribbon with the superimposed models coloured red, green, orange, blue and pale gold and represented as ribbons. Panel C depicts the two main strategies for the sugar targets. The top figure concerns target 130 and shows the restraints extracted from structure (among others) 2uvj. The receptor is shown as blue cartoons.

with the binding site residues depicted as lines too. The sugar is shown as sticks with the carbon and oxygen atoms being coloured dark green and red respectively. The bottom figure concerns the remaining sugar targets and shows our top-ranking model for target 128 (medium quality). The consensus binding site that was defined based on the available templates is coloured orange, whereas the rest of the receptor is shown as transparent surface and cartoon and coloured blue.

For the scoring experiment, we first identify the largest meaningful common subset of residue contacts across all models as a way of standardising the set on which we score. That sometimes involves excluding models which are not docked or have very small/peculiar interfaces. Then, all missing atoms are rebuilt according to OPLS topologies. Finally, all models underwent a short energy minimisation (50 steps of steepest descent EM) and are scored according to the *HS_itw_* function shown above (minus the restraint term) when dealing with protein-protein systems; The scoring term weights are adjusted accordingly for different systems (see below for details).

### Targets involving peptides

#### Target 121 (round 38)

We modelled the protein (TolA) on PDB entry 1lr0, renumbering the residues as required to match the target sequence. For the peptide we created an ensemble of 3 models: The first and second models were created in PyMOL^31^ using the known sequence of the peptide and ideal backbone angles that corresponded to beta and polyproline-II conformations; the third was obtained from the PEPFOLD webserver^32^.

For the server submission we performed ensemble docking with increased sampling using restraints we identified in the literature^33^ to drive the docking. For the manual submission we performed the joint analysis mentioned previously (see second to last paragraph of the *Methods* section) over a plethora of runs, which centred around the concept of specifying restraints that would mimic hydrogen bonds between the peptide and the protein.

#### Targets 134 and 135 (round 44)

For the server submission of target 134, we modelled the protein on PDB entry 4d07 and the peptide on 4d07 and 4qh8, threading the target sequence on the peptide structure of the templates. For the manual submission, we defined restraints to dock instead of threading and refining.

For target 135, we used the same templates for the peptide as target 134 (4d07 & 4qh8) and added 5el0 and followed similar strategies as for target 134.

### Template-based targets

#### Target 122 (round 39)

We modelled the IL23R receptor using MODELLER and the RAPTOR-X webserver creating an ensemble of two models, IL12B using FATCAT^34,35^ in addition to the model that was provided by the organisers. For IL23A we used the unbound structure that was provided by the organisers.

For the server submission we superimposed the models of IL12B and IL23A onto PDB structure 5e4e and then, using the IL23A subunit, which is shared between PDB entries 5e4e and 2d9q, we superimposed all models onto 2d9q to create a model of the entire complex which was then refined in HADDOCK. For the manual submission, we extracted interface Cα-Cα restraints^18^ from multiple templates and used those to drive the docking instead of superimposing and refining.

#### Target 136 (round 36)

Our main strategy for this target was to model the monomers based on available templates and then recreate the full decamer using symmetry restraints. For the manual round, we extracted interface information from the top model (according to the HADDOCK score) of the server submission and turned them into restraints to drive the docking. These runs were performed using our new MARTINI^36,37^ coarse grain implementation^38^.

### Information-driven targets

#### Targets 123 and 124(round 39)

For the server submission of target 123, we generated an ensemble of models for the PorM protein using the Raptor-X and Robetta^39^ webservers and an ensemble of models for the nanobody using MODELLER and PDB entries 5imo and 4qlr as templates. We docked using the CDR residues of the nanobody as active residues and the surface residues of PorM as passive residues with increased sampling (10000/400/400 instead of the default 1000/200/200 models). For the manual submission we repeated the same run, but we replaced the nanobody ensemble with one kindly provided to us by Dr. Jeff Gray (Johns Hopkins Medical School, Baltimore) (based on PDB entry 5lmw). We submitted the top model of the top 10 clusters for the server and the top 10 models (irrespective of clustering) for the manual prediction.

We could not identify any usable information for target 124, either in the form of template structures or interaction data. Instead we defined custom restraints in order to generate models with the global shape of a H (for Haddock, or Hopeless) in order to provide some unrealistic but well optimised interfaces for scoring.

#### Target 125 (round 40)

For the server round, we used available structures for the LLT1 dimer (PDB entries 4qkh, 4qki, 4qkj and 4wco) after renumbering them so they matched the target numbering to create a dimer ensemble. We modelled the NKR-P1 dimer on PDB entries 3m9z and 3t3a using MODELLER to cover all possible combinations of types of interfaces and interacting loops that were available in the literature^40,41^. We drove the docking using interface information extracted from the available complexes^42–44^ and the literature^40,41,45^. For the manual submission we repeated the same runs as for the server submission but with a filtered list of residues.

#### Targets 131 and 132 (round 42)

A structure of CEACAM1 available in the PDB (entry 2gk2) was used for the docking of both targets. For the modelling of HopQ1, we used template chain A of PDB entry 5lp2 which has an almost perfect sequence identity with our target sequence but does not cover the entire target sequence. To model those gaps we used MODELLER and selected the top 10 models based on the DOPE score. For HopQ2, we created an ensemble of three models, in which the first came from the RAPTOR-X webserver, the second from iTasser and the last was created in MODELLER after manually curating the alignment between our target sequence and chain A of 5lp2. For the docking we made use of information available in the literature^46^, which suggested that a beta strand of HopQ is crucial for the CEACAM1-HopQ interaction and two mutations of CEACAM1 almost knocked out the interaction altogether.

#### Target 133 (round 43)

For the server round, to drive the docking, we mapped the sequence differences between target and template proteins to the sequence of the target and defined those restraints as active. For the manual round, we performed a plethora of runs, trying different ways of specifying the restraints (with or without random removal), using single models instead of ensembles, using ROSETTA and molecular dynamics (with GROMACS^47,48^) to sample alternative conformations of the proteins.

### Targets involving glycans

#### Targets 126-130 (round 41)

We modelled the receptors with ROSETTA-CM protocol^49^ using the templates 2xd3, 2xd3, 3k00, 2uvj for targets 126-129, respectively and template 3cu9 for target 130. In addition to the already mentioned templates, we identified many more with HHPRED using the sequence of the target and filtering for the presence of saccharides in the binding pocket. We used those templates to define both interfaces and more specific ambiguous restraints between the sugar molecules and the receptors. For the manual round, we further refined the list of templates that we were using as well as expand the selection of residues specified as active. No other settings were changed compared to the server submission.

For the ligands, we generated up to 500 conformers using OpenEye OMEGA^50,51^, clustered them and selected representative structures that resulted in the creation of ensembles with 6, 9, 6, 4 and 4 conformations for targets 126-130, respectively.

## Results and discussion

Table 1 summarises the results we obtained for the server and manual predictions and for the scoring experiments. In total, we generated acceptable-or higher-quality models for 13 and 11 interfaces for the server and manual submissions, respectively, which corresponds to success rates of 59 and 50% when considering the top 10 models submitted for evaluation. If we consider targets instead of interfaces, and consider successful any target for which at least one interface was correctly predicted, then our success rates jump to 63% and 56%, respectively. Our performance in the scoring experiment is even better, with 14 (63%) interfaces and 11 (69%) targets successfully predicted.

**Table 1:** Summary of the prediction and scoring results obtained by the HADDOCK server and manual team. The first column is the target and interface number as reported by CAPRI and the second through fourth columns refer to the number of high-(***)/medium-(**)/acceptable-(*)quality models generated during the server, manual and scoring experiments, respectively, when considering the top ten models submitted for evaluation.

| Target | Target Category | Server<br>***/**/* | Manual<br>***/**/* | Scoring<br>***/**/* |
| --- | --- | --- | --- | --- |
| 121 | Peptide | 0/0/0 | 0/0/0 | 0/0/0 |
| 122 | Template-based | 0/0/0 | 0/0/0 | 0/1/0 |
| 123 | Information-driven | 0/0/0 | 0/0/0 | 0/0/0 |
| 124.1 | Information-driven | 0/0/0 | 0/0/0 | 0/0/0 |
| 124.2 |  | 0/0/0 | 0/0/0 | 0/0/0 |
| 125.1 | Information-driven | 0/0/4 | 0/0/0 | 0/4/0 |
| 125.2 |  | 10/0/0 | 7/1/2 | 7/3/0 |
| 125.3 |  | 0/0/0 | 0/0/0 | 0/0/0 |
| 125.4 |  | 0/0/0 | 0/0/0 | 0/0/0 |
| 126 | Glycans | 0/0/1 | 0/0/0 | 0/0/3 |
| 127 |  | 0/1/1 | 0/0/4 | 0/0/5 |
| 128 |  | 0/0/10 | 0/2/7 | 0/2/8 |
| 129 |  | 0/0/5 | 0/0/6 | 0/0/7 |
| 130 |  | 1/5/4 | 2/4/3 | 1/5/2 |
| 131 | Information-driven | 0/0/0 | 0/0/0 | 0/0/0 |
| 132 | Information-driven | 0/0/0 | 0/0/0 | 0/0/1 |
| 133 | Information-driven | 0/3/3 | 0/3/3 | 0/1/5 |
| 134 | Peptide | 0/0/1 | 0/1/1 | 0/2/0 |
| 135 | Peptide | 0/0/5 | 0/4/1 | 0/0/0 |
| 136.1 | Template-based | 0/5/5 | 0/9/1 | 0/4/5 |
| 136.2 |  | 0/4/1 | 0/0/8 | 0/1/3 |
| 136.3 |  | 0/0/8 | 0/0/1 | 0/1/5 |

Breaking down our performance by target category, we generated medium-and acceptable quality models for two (targets 134 and 135) of the three peptide targets (targets 121, 134 and 135), with target 121 being a challenging target for which only 6 acceptable or higher quality models could be found among all submissions. For target 134 we correctly predicted the binding 12mer sequence out of the 50-residue long peptide for both manual and server submissions making good use of the information available in the literature, but failed to generate higher quality models for target 135 for which the binding residues were known, although we did generate more models of the same quality in the top 10. Targets 134 and 135 serve as a good example of the difficulty we are facing in scoring good models in the top 10 as can be seen in Table S1, where for the manual submission of target 135 we only generated 4 medium-and 1 acceptable-quality model but when considering 43 models instead of ten we are generating 11/23/4 high-/medium-/acceptable-quality models instead. This pattern holds true for almost all targets for which more than the top 10 models were evaluated indicating there is plenty of room for improvement in the scoring of our models.

Despite the fact targets 122 and 136 were template-based, we only achieved good performance for the latter, generating acceptable models for all three interfaces and medium quality ones for two of the three for the server submission. This target is particularly interesting as its structure was determined with cryo-EM and it is the largest complex featured so far in CAPRI. Therefore, it was an excellent test case for our implementation of the MARTINI coarse grain forcefield into HADDOCK^38^. Using less than ideal information (interface restraints extracted from the top model of the all-atom, server submission) we were able to accurately recreate the full decamer using five dimers with interface restraints specified only between the first two adjacent ones (i.e. simultaneous five-body docking) and symmetry restraints to recreate the rest of the complex, while at the same time massively reducing the computational cost from approximately 10-12 hours to 2 hours per model (5-6 fold speedup).

For the majority of the targets we followed an information-driven strategy as we were able to identify some information about the putative interfaces of the target complexes that could aid the modelling process. Unfortunately, the information we uncovered was not enough to overcome the challenging nature of most of these targets (1 acceptable-quality model for target 123, no acceptable-quality models for 124, 1 medium-quality model for 131 and 1 medium-and 4 acceptable-quality models for 132 when considering the top 10 models). We successfully modelled 2 of the 4 interfaces of target 125 (1 and 2), generating acceptable quality models for the first as well as high and medium quality for the second. We were not able to generate acceptable models for the other two interfaces which proved too challenging for all the participants as only one acceptable model was generated for the third interface and none for the fourth. We were able to generate good models for the last of the information-driven targets, 133, achieving the same performance for server and manual predictions. Preliminary analysis of these targets (the crystal structures were not yet available at the time of writing) indicates that the limiting factor for these targets was poor modelling of the individual components of the complex rather than the docking itself.

The last targets were those of round 41 which featured glycans. These targets also represent a first, as they were the first protein-small molecule complexes to be featured in CAPRI and HADDOCK performed well across all of them, generating acceptable models for all but the manual submission of target 126, and even generating highly accurate models for both the manual and server submission of target 130. The topologies and parameters for describing the glycans were automatically obtained from PRODRG^52^, after modifying their numbering and naming to make them look as one single heteroatom residue.

Our scoring performance ranks us as one of the best performers of the scoring experiment when considering the number of targets for which acceptable models were identified with acceptable-or higher quality models in 14 of 22 interfaces. The robustness of our scoring protocol and function is also demonstrated in targets such as 132 for which we were able to pick out one of the 27 acceptable models in a pool of over 2000 models as well as the fact we could identify near-native models in all target categories. One limitation of our scoring protocol, which also affects our CAPRI ranking, is the difficulty in distinguishing medium-or higher quality models from acceptable-quality ones.

## Conclusions

CAPRI rounds 38-45 featured many firsts such as the inclusion of a cryo-EM determined target in target 136 of round 45 and protein-small molecule (glycans) complexes in targets 126-120 of round 41. HADDOCK was able to generate near-native models for these systems as well as most of the traditional protein-protein systems that featured in the remaining targets, including the peptide ones. While it is evidently clear the entire docking community still has difficulties in dealing with systems for which no good templates are available, this is an issue that seems to be affecting our efforts in CAPRI for the remaining targets as well: As it appears poor initial modelling of the components rather than the docking itself is affecting our performance. The second limiting factor is our scoring performance during the docking experiments (manual and server). As can be seen in Table S1, for every target for which more than the top 10 models have been evaluated, there are many higher quality models that we are failing to rank in the top 10. This creates a contradiction with our performance in the scoring experiment for which we can reliably identify near-native models in a pool of thousands and indicates that there is room to re-optimise our scoring function. Development and testing of the next major version of HADDOCK (v2.4) is currently ongoing and we expect that many new features and small improvements will allow us to both generate higher quality models as well as rank them more accurately.

## Acknowledgements

The authors would like to thank Dr Cristina Paissoni for contributions she made for target 133.

