## Supplemental Information for "An overview of data-driven HADDOCK strategies in CAPRI rounds 38-45"

### Detailed descriptions of the docking strategies per target

#### Targets involving peptides

##### Target 121

This was a protein-peptide complex determined by NMR consisting of the C-terminal domain of TolA and a small peptide, likely part of the N-terminal region of TolB.

For the server submission, we performed ensemble docking with sampling setting of 6000/400/400 it0, it1 and itw models, respectively, full flexibility and backbone dihedral angle restraints for the peptide. To drive the docking, we used protein residues (24, 25, 34, 35, 36, 37, 39, 40, 48, 56, 65, 69, 94, 96, 97, 98) and all residues of the peptide as active. These residues were identified in a series of NMR titration experiments of TolA with the N-terminal domain of TolB. We used all protein residues which showed chemical shift perturbations at molecular equivalents of peptide concentration <= 1. We submitted the top models of the top cluster and the top 1 model of all subsequent clusters to round out the top 10.

For the manual submission we performed the joint analysis mentioned in the *Methods* section (second to last paragraph). For these runs we set a positive charge to the peptide N-terminus, and followed our protein-peptide docking protocol[1] (doubling it1 steps, setting the number of it1 structures to be analysed to 400, using RMSD with a cut-off value of 5Å for the clustering and disabling secondary structure dihedral angle restraints).

##### Targets 134 and 135

These targets featured the binding of a small (12meric) peptide to the homodimeric (A2) structure of Dynein Light Chain subunit 8 (DLC8) in two distinct experiments. For the first, the complex structure was solved using a 50-residue long peptide that included the aforementioned 12meric peptide and for the second only the 12meric peptide was cocrystallised with DLC8. In both experiments, the same binding pose was adopted by the peptide resulting in the formation of the same complex. The challenge of target 134 centred around predicting which 12mer of the 50-residue long peptide was the part that bound to DLC8. For target 135 the 12 binding residues of the peptide were circulated to the participants; The challenge of this target was to produce highly accurate models of the protein-peptide complex.

For the protein we used the homodimeric form of the receptor and for the peptides we threaded the entire sequence of the 50-residue long peptide in 12-residue long slices on the respective peptides of 4d07 and 4qh8 for a total of 39 independent refinements. All mutations and rebuilding of mutated sidechains were handled by HADDOCK, after sampling multiple side chain conformations. Additionally, we applied C2 and NCS restraints between the two symmetric protein and peptide chains. We submitted the top model of the top cluster of each docking run with the two peptide templates mentioned previously.

For the manual submission, we defined hydrogen-bond restraints that would place the peptide in antiparallel conformation relative to the β-strand of the receptor against which it binds, disabled the symmetry restraints and used RMSD values for clustering with a cut-off of 5Å, and selected models for submission based on visual inspection and the presence of sequence motifs[2,3] which were suggested as crucial for the binding to take place.

For target 135, we extended the structure of our picks for the best template from target 134 (the peptide with residue index 29 out of the 50-residue long peptide) by one residue (L) to match the target sequence. Similarly, to target 134, we then defined hydrogen-bond restraints between peptide and receptor and docked, submitting the top 10 models according to the HADDOCK score.

For the manual submission, we analysed a combination of runs consisting of the hydrogen-bond-driven ones mentioned above and refinements similar to the ones mentioned in the server section of target 134 but with the desired target sequence peptide alone. We submitted the top 4 models of the top cluster and the top model of all subsequent clusters from the combined analysis.

#### Template-based targets

##### Target 122

Target 122 was a human cytokine receptor complex (IL23-IL23R), consisting of the receptor IL23R bound to cytokine heterodimer IL23, itself comprised of subunits IL23A and IL12B for a total of three chains with a stoichiometry of 1:1:1.

##### Target 136

Target 136 was the first CAPRI target whose structure has been determined by cryo-EM instead of X-RAY crystallography or NMR. It was a homodecameric lysine decarboxylase which adopted D5 symmetry. We based our modelling on PDB entries 2vyc and 3n75.

For the server submission, we first generated models of the monomers of our target using the two templates separately. We then superimposed the monomer ensembles on their respective templates to generate the dimers along the C2 symmetry axis and refined them. The resulting models were clustered and the top scoring models of the top 5 clusters were selected for an ensemble in which the first 5 models were based on 2vyc and the latter 5 on 3n75. This ensemble of dimeric models was then superimposed on the 5 dimers of the 2vyc template and refined again to recreate the full decamer using C5 symmetry restraints. Due to the size of the system, we disabled water refinement and instead only performed a short EM. We also disabled cross-docking for all runs.

For the manual submission, we identified the top model of the server submission that originated from 2vyc and extracted interface restraints from it along the C2 axis using a distance cut-off of 3.5Å. Those restraints were used to drive four docking runs in total (two per template), in the first of which the flexible regions were defined by HADDOCK automatically (default behaviour) and in the second we defined the hinge regions between the domains of the chains as fully flexible – meaning they would have full flexibility during all it1 stages. For these runs we coarse-grained the top 5 monomer models of both templates according to the MARTINI 2.2[4,5] force field HADDOCK implementation[6], used the interface restraints mentioned above, disabled cross-docking, disabled the systematic sampling of 180° rotated solutions during it0, skipped it0 and set the number of models per stage to 5000/400/400, respectively. We submitted the top 5 models of the 2vyc-based run without flexibility, the top 4 models of the 2vyc-based run with flexibility and the top models of 3n75-based run with flexibility.

#### Information-driven targets

##### Targets 123 and 124

Targets 123 and 124 focused on the PorM protein. In the former, the target was the N-terminal domain of PorM complexed with a nanobody and in the latter the dimeric C-terminal PorM domain complexed with a different nanobody. Both targets were very challenging with only one acceptable model when considering all 100 submitted models for Target 123 and no acceptable models for Target 124.

##### Target 125

Target 125 was a human hetero hexamer made up of the extracellular domain of LLT1 and the extracellular domain of its inhibitor, NKR-P1.

For the server submission, residues 97, 99, 101, 112 and 113 and their symmetry pairs, and 97, 100, 101, 102, 112, 113, 114, 117 and 118 and their symmetry pairs were specified as active for LLT1 and NKR-P1, respectively. We also used C2 and NCS restraints for the NKR-P1 homodimers and increased the sampling to 2000/200/200 for the three stages, respectively. We submitted the top three models of the top two clusters followed by the top 2 models of the next two clusters.

For the manual submission, we filtered the residues that were used during the server round based on the mean solvent-accessible surface area of the starting ensembles and repeated all docking runs. We then analysed all models and submitted the top three models of the top cluster, the top two models of the next three clusters and the top model of the fifth-best cluster following the combined analysis.

#### Targets 131 and 132

Targets 131 and 132 featured the human CEACAM1 protein in complexes with the cell adhesion proteins HopQ type I and HopQ type II, respectively. This was another challenging target with only one and four acceptable models in the top 10 by one and two submitting groups respectively; More medium- and acceptable-quality models were available in the top 100 set.

For target 131, residues 91, 92, 94, 98, 99, 100, 101, 102, 103, 104, 105, 106, 108, 110, 170, 172, 174, 176, 179, 180, 181, 182, 183, 184, 186, 187, 188, 189, 190, 191, 192, 194, 196 of HopQ1 were defined as passive and residues 35 and 92 of CEACAM1 as active. The sampling was increased to 5000/400/400 for the three stages respectively and we also disabled the random removal of restraints owing to the low number of active residues. For the server round, we submitted the top four models of the top cluster and the top model of all subsequent clusters. For the manual submission, we analysed all runs that were performed during the server round and submitted a representative selection of the top clusters.

For target 132, residues 96, 98, 111, 112, 113, 122, 123, 124, 125, 126, 162, 169, 181, 182, 183, 184, 185, 186, 187, 188, 189, 190, 192, 197, 200, 202 of HopQ2 were defined as passive and residues 35 and 92 of CEACAM1 as active. The sampling was increased to 6000/400/400 for the three stages respectively and we also disabled the random removal of restraints. For the server round, we submitted the top four models of the top cluster and the top model of all subsequent clusters. For the manual submission, we analysed all runs that were performed during the server round and submitted a representative selection of the top clusters.

#### Target 133

Target 133 featured a redesigned version of the Colicin E2 DNAse-Im2 complex, with the redesigned components being labelled $E^{des3}$ and $Im^{des3}$. We modelled both chains on the respective chains of the wild-type complex (PDB entry 3u43) using MODELLER and creating an ensemble of 10 models for both partners after ranking them by the DOPE score.

For the server submission residues 59, 73, 76, 77, 80, 83, 89, 90, 97, 98, 99 of $E^{des3}$ and residues 21, 23, 25, 30, 39, 42, 59 of $Im^{des3}$, respectively, were defined as active. We also increased the sampling to 10000/400/400 for the three stages respectively. We submitted the top model of the top 10 clusters.

For the manual submission, we analysed all models of all runs and selected representative structures of the top clusters. The top 5 models do not deviate heavily from the wild-type complex, whereas the next 5 sample more freely.

### Targets involving glycans

#### Targets 126-130

CAPRI round 41 was the first time that protein-small-molecule complexes were the focus of the competition. Specifically, for targets 126-129 the challenge was to predict the binding pose of arabino-saccharide ligands of varying lengths (3-6 copies of the sugar molecule) complexed with a binding protein which is part of a larger ABC transporter which transfers arabino-oligosaccharides into the cell. For target 130 we had to predict the complex structure of one of the sugars (the pentose), bound to a mutated version (E201A) of the AbnB protein.

For the server round, the residues specified as active were 14, 45, 46, 47, 48, 51, 69, 70, 72, 130, 184, 186, 236, 237, 238, 239, 240, 259, 288, 292, 327, 376, 377 for targets 126 and 127, residues 14, 15, 16, 19, 46, 47, 50, 70, 71, 72, 130, 184, 185, 186, 259, 288, 290, 291, 292, 373, 376 for target 128 and residues 14, 15, 46, 69, 130, 179, 181, 182, 239, 240, 259, 260, 264, 291, 292, 376 for target 129. The latter targets feature the smallest saccharides (hexose to triose) which is why their respective restraints are not as extensive. The restraints defined mimic the hydrogen bonds formed between the ligand and their respective templates and were specified from the active residues listed above to any oxygen atom of the ligand, with a target distance appropriate for hydrogen bonds. For target 130 restraints were defined between protein residues 26, 146 and 200, and specific sugar oxygens and were not subjected to random removal. The restraints for target 130 were extracted from available templates as well as relevant literature[7]. We also lowered the scaling of intermolecular interactions to a tenth and a thousandth of its original value, for targets 126-129 and 130, respectively, to allow the ligands to overcome the van der Waals barrier and more easily penetrate into the binding pocket. We adjusted the weight of the vdW energy during the scoring of it0 to a tenth of its original value so as to not penalise the clashes that would inevitably arise from using a lowered intermolecular scaling constant. We used RMSD clustering with cut-off values of 4, 3.5, 3.5, 3 and 3.5Å for targets 126-130, respectively. We increased the sampling during it0 to 10000 models for targets 126-129 and 400 for target 130 while the number of models generated during the flexible stages for all models was set to 400. All the ligand molecules were specified as fully flexible.

***Table S1:*** *Summary of the results obtained by the HADDOCK team for the manual, server and scoring experiments. The first column is the target and interface number as reported by CAPRI, the second and third columns refer to the number of high-/medium-/acceptable-quality models generated during the manual experiment, when considering the top 10 and 100 models submitted for evaluation, respectively and the fourth and fifth columns refer to the number of high-/medium-/acceptable-quality models generated during the server experiment, when considering the top 10 and 100 models submitted for evaluation, respectively. * marks targets for which more than the top 10 models were evaluated. Targets highlighted in bold indicate successful submissions.*

| **Target** | **Manual top 10** ***/**/* | **Manual top 100** ***/**/* | **Server top 10** ***/**/* | **Server top 100** ***/**/* |
|---|---|---|---|---|
| 121 | 0/0/0 | 0/0/0 | 0/0/0 | 0/0/0 |
| 122 | 0/0/0 | 0/0/0 | 0/0/0 | 0/0/0 |
| 123 | 0/0/0 | 0/0/0 | 0/0/0 | 0/0/0 |
| 124.1 | 0/0/0 | 0/0/0 | 0/0/0 | 0/0/0 |
| 124.2 | 0/0/0 | 0/0/0 | 0/0/0 | 0/0/0 |
| **125.1*** | **0/0/0** | **0/0/12** | **0/0/4** | **0/0/36** |
| **125.2*** | **7/1/2** | **44/41/15** | **10/0/0** | **64/28/8** |
| **125.3*** | **0/0/0** | **0/0/0** | **0/0/0** | **0/0/1** |
| 125.4 | 0/0/0 | 0/0/0 | 0/0/0 | 0/0/0 |
| **126*** | **0/0/0** | **0/0/2** | **0/0/1** | **0/0/13** |
| **127*** | **0/0/4** | **0/0/30** | **0/1/1** | **0/1/18** |
| **128*** | **0/2/7** | **0/13/78** | **0/0/10** | **0/3/84** |
| **129*** | **0/0/6** | **0/0/49** | **0/0/5** | **0/2/40** |
| **130*** | **2/4/3** | **6/74/19** | **1/5/4** | **2/46/52** |
| 131 | 0/0/0 | 0/0/0 | 0/0/0 | 0/0/0 |
| **132*** | **0/0/0** | **0/0/9** | **0/0/0** | **0/0/0** |
| **133*** | **0/3/3** | **0/9/33** | **0/3/3** | **0/24/39** |
| **134*** | **0/1/1** | **0/1/5** | **0/0/1** | **0/3/14** |
| **135*** | **0/4/1** | **11/23/4** | **0/0/5** | **0/0/5** |
| 136.1 | 0/9/1 | 0/9/1 | 0/5/5 | 0/5/5 |
| 136.2 | 0/0/8 | 0/0/8 | 0/4/1 | 0/4/1 |
| 136.3 | 0/0/1 | 0/0/1 | 0/0/8 | 0/0/8 |
